# Radial-axial transport coordination enhances sugar translocation in the phloem vasculature of plants

**DOI:** 10.1101/2021.09.24.461704

**Authors:** Mazen Nakad, Jean-Christophe Domec, Sanna Sevanto, Gabriel Katul

**Author notes:** **Author contributions:** G.K. conceived the original screening and research plans, JC.D., S.S. and G.K. supervised the progress of the model development and numerical analysis, M.N. conceived the project and wrote the article with contributions from all the authors, M.N. agreed to serve as the author responsible for contact and communication. **Corresponding author:** Mazen Nakad.

## Abstract

Mass transport of photosynthates in the phloem of plants is necessary for describing plant carbon allocation, productivity, and responses to water and thermal stress. Several hypotheses about optimization of phloem structure and function and limitations of phloem transport under drought have been proposed, and tested with models and anatomical data. However, the true impact of radial water exchange of phloem conduits with their surroundings on mass transport of photosynthates has not been addressed. Here, the physics of the Munch mechanism of sugar transport is re-evaluated to include local variations in viscosity resulting from the radial water exchange in two dimensions (axial and radial). Model results show that radial water exchange pushes sucrose away from conduit walls thereby reducing wall frictional stress due to a decrease in sap viscosity and an increase in sugar concentration in the central region of the conduit. These two co-occurring effects lead to increased sugar front speed and axial mass transport across a wide range of phloem conduit lengths. Thus, sugar transport operates more efficiently than predicted by previous models that ignore these two effects. A faster front speed leads to higher phloem resiliency under drought because more sugar can be transported with a smaller pressure gradient.

**Summary:** The overall speed of sap increased by including a concentrationdependent viscosity in axial and radial directions.

## 1 Introduction

The efficiency of photosynthate transport from the production sites (sources; usually leaves) to areas of consumption or storage (sinks) within the vascular tissue known as the phloem is drawing significant attention in plant physiology. The implications of efficient photosynthate transport range from local impacts on tissue or plant health and growth to ecosystem-scale effects on carbon and water cycling because of the potential link between phloem transport and stomatal control of photosynthesis (Nikinmaa et al. 2013), and a possible link between phloem transport failure and plant mortality under drought (Sevanto et al. 2014). Consequently, several models for phloem transport and the connection between phloem structure and function, as well as for the potential weak points in the transport system have been formulated (Munch 1930, Phillips & Dungan 1993, Thompson & Holbrook 2003, Jensen et al. 2009, 2012, Sevanto 2014). The most commonly accepted concept under which all these models operate is that phloem vasculature is optimized for efficient transport of soluble organic compounds (mostly sugars) produced during photosynthesis approximately as described by the pressure-flow hypothesis or Münch mechanism (Münch 1930). In the pressure-flow hypothesis, transport is initiated in leaves when sugars and other metabolic products are loaded into the phloem. Once in the phloem, sugars and water molecules are driven to move through the phloem’s complex network of narrow but elongated, interconnected, cylindrical living cells (sieve tubes) spanning the length of the plant following pressure gradients. High sugar concentration at the loading site (leaves) draws water from the xylem, the tissue supplying water to the leaves or other surrounding tissues by osmosis towards the phloem. At the sinks, sugars are unloaded from the phloem to growing or storage cells, and water is released back to the xylem or other surrounding tissues. The loading and unloading at sources and sinks build a pressure gradient in the phloem creating a system where plant water and photosynthate transport over long distances occurs without active pumping. This cycle of pressure buildup and transport without active pumping endowed the phloem system with the label *miracle of ingenuity* (van Bel 2003).

The simplicity, plausibility, and intuitive appeal of the Munch mechanism lead to its proliferation in mathematical models (Nikinmaa et al. 2013, Jensen et al. 2016). It is routinely used to connect plant carbon sources and sinks, and their concomitant controls in a future CO_2_ enriched climate (Mencuccini & Hölttä 2010, Fatichi et al. 2019), and it has been used to explain aspects of plant hydraulic failure during drought (Savage et al. 2017, Konrad et al. 2018, Huang et al. 2018, Sevanto 2018, Salmon et al. 2019) and extreme cold temperatures (Swanson & Geiger 1967, Wardlaw 1968). The direct consequence of those two abiotic stresses should be a decrease in the overall phloem flow rate because the viscosity of a sucrose solution increases significantly with the drought-induced increase in sugar concentration required for osmoregulation (Hölttä et al. 2009) and with decreasing temperature.

However, the Münch mechanism is not free from controversy. The main critique stems from the fact that the sieve tubes seem to have too low of a hydraulic conductance along the phloem to allow sugars to be transported from leaves to roots in the largest and longest of plants (Curtis & Scofield 1933, Lang 1979, Fensom 1981, Knoblauch et al. 2016, Liesche & Schulz 2018). These studies also report lower leaf sucrose concentration in tall trees compared to shorter vegetation. When interpreted using simplified transport models for hydraulic conductance, this suggests that the Münch mechanism cannot produce effective transport in tall plants because the driving force for sucrose transport is lower for a longer path length. To resolve the issues of inadequate conductance and too low-pressure gradient to efficiently drive flow in tall plants, it has been suggested that rather than exchanging water and sugars only at the extreme source and sink ends of the phloem pathway, as suggested by the original Munch mechanism, sugars and water could be exchanged at different locations along the pathway essentially forming a ‘‘relay” system to facilitate transport (Lang 1979). While plausible, and based on modeling studies, also effective in increasing transport capacity (Hölttä et al. 2009), there currently is no clear evidence for unloading and reloading of sugars from and to phloem conduits along the transport pathway. There is, however, an increasing amount of evidence that water might readily be exchanged between the conduits and their surroundings along the entire length of the pathway (Knoblauch & Oparka 2012, Knoblauch & Peters 2017, Stanfield et al. 2017).

Answering the question of whether and how easily phloem conduits exchange water with their surroundings outside the primary loading and unloading zones at sources and sinks is becoming necessary for evaluating the validity of the Münch mechanism, and because it determines how phloem transport is affected under stress (Sevanto 2014, 2018). Theoretically, if no water exchange occurs, plants run a risk of blocking phloem flow by viscosity increase and reduced pressure gradient under drought conditions because higher amounts of sugar are needed in the phloem conduits at the loading and unloading zones for osmoregulation against the declining water potential of the xylem and the surrounding tissues. If water exchange occurs readily along the entire transport pathway, the flow may not be restricted by the same constraints that stem from the original interpretation of the Munch mechanism (Phillips & Dungan 1993, Sevanto 2014, Sevanto et al. 2014, Sevanto 2018). In particular, the effects of viscosity buildup can be ameliorated because of the diluting effects of radial water flow, the focus of the work here.

Experimental challenges in measuring water and solute fluxes within the phloem (Curtis & Scofield 1933, Housley & Fisher 1977) have led to reliance on mathematical models of simplified phloem transport to understand transport mechanisms in the phloem. As expected when employing such models, values of one or more variables may not be well constrained or are uncertain, possibly by several orders of magnitude. This fact is often used to justify (overly) simplified description of transport physics in models. These simplifications might lead to biased mass fluxes and estimates for total transport. An alternative to the simplification approach is to assess the effects of the simplification on the results, and in the case of phloem transport, test whether the pressure-flow hypothesis predicts increases or decreases in sugar mass flux when these simplifications are relaxed or re-addressed. If increases in mass flux can be demonstrated upon re-addressing key simplifications used with the Münch mechanism, the contradictions with some observations might be explained by the effects of these model simplifications lending further support to the Münch mechanism.

Irrespective of the physics of phloem transport, phloem anatomical structure is assumed to have evolved to optimize sugar transport. Within the confines of the Münch mechanism, this optimization arises because of trade-offs between benefits of increasing sugar concentration (*c*) and its impact on the mass flux (*J*). Increasing *c* increases flux *J* because the osmotic pressure driving water movement increases approximately linearly with *c* (i.e. Van’t Hoff equation); however, increasing *c* is accompanied by a nonlinear increase in dynamic (and kinematic) viscosity thereby enhancing the viscous forces that oppose movement (drag) thereby reducing *J*. In most phloem transport models today, viscosity is treated either as a constant or it is allowed to vary with loaded sugar concentration assuming that radial water flow does not significantly affect sugar concentration or viscosity. Theoretical representation of *J* along with a number of scaling arguments results in a maximal sugar flux *J_max_* at around *c* = 20% wt/wt (Jensen et al. 2013) independent of the sieve tube geometric properties. Interestingly, upon averaging across species and experiments, the operating *c* = 20% wt/wt was reasonably confirmed and appears independent of properties of the sieve tube geometry or the loading mechanism (passive versus active) in plants. The scatter in reported values of *c* around *c* = 20% wt/wt, however, was substantial (Jensen et al. 2013) with many species operating at *c* < 20% wt/wt. This low loaded sugar concentration value has also been used to argue against the validity of the Munch mechanism, especially in tall trees (Knoblauch & Oparka 2012), since it leads to a decrease in mass flux as predicted by the simplified physics. Therefore, it remains open whether plant sugar transport actually operates sub-optimally, or whether alternatives or modifications to the Münch mechanism are necessary to explain long-distance yet sub-optimal sugar transport.

Motivated by these issues, we ask to what degree refinements and addressing the model simplifications in the description of the transport physics within the Münch mechanism enhance *J* above and beyond expectations from earlier theories. To address this question generically, an idealized, unsteady, two-dimensional, osmotically driven pipe flow governed by the physics of the Münch hypothesis is considered. No attempt is made to represent all the complexities of the geometry of the phloem tissue or in the loading and unloading mechanisms of sugars. Instead, the main novelty here stems from the inclusion of the simultaneous effects of concentration-dependent viscosity where local changes in viscosity with *c* (axially and radially) are allowed. It is shown that including such adjustments to viscous stresses lead to significant enhancement in the magnitude of the mass flux, especially in long tubes. Moreover, this enhancement in *J* is shown to be accompanied by a reduced pressure gradient driving the flow. Thus, the work here adds support to the Munch hypothesis by offering a new perspective regarding the contribution of coordination between axial and radial flow to *J*. This coordination is enhanced especially in long tubes which explains why high sugar concentration is not needed in tall tree.

Before presenting the results of this new representation of viscous stresses, some comments and clarifications about efficient sugar transport and its relation to an optimal *c* are illustrated through the occurrence of a maximum sugar flux *J* = *J_max_* at a well-defined *c* in steady-state and globally averaged Poiseuille flow models (Jensen et al. 2013). Measured sugar concentration in the leaves of tall trees being lower than this optimal *c* corresponding to maximum sugar flux was the basis for some critique of the Munch hypothesis (Knoblauch et al. 2016). It is to be noted that the c corresponding to *J_max_* in steady-state and globally averaged Poiseuille models is shown not to be sensitive to the phloem hydraulic properties or even tube geometry. Hence, the occurrence of such a *c* is weakly connected to phloem hydraulics as later discussed.

In prior work, a tube of constant length *L* and radius *a* was considered with *a/L* ≪ 1. The sugar mass flux *J* (kg s ^-1^) was assumed to be only advective and given by

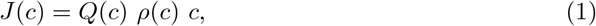

where *Q*(*c*) is the volumetric flow rate (m^3^ s^-1^), *ρ*(*c*) is the density of the phloem sap that varies with *c*, and *c* is the sucrose concentration inside the sieve tube as before (Jensen et al. 2013). In this approach, the driving force for *Q* and the constraints on it are now formulated to be *c* dependent. The maximal flux *J_max_* emerges when solving for c at the critical point *∂J/∂c* = 0. For *c* > 0, the existence of this single critical point is virtually guaranteed provided the water flux *ρ*(*c*)*Q*(*c*) non-linearly decreases with increasing *c*. For laminar flows in tubes, the Hagen-Poiseuille (HP) equation for *Q* and the resulting *J* can be expressed as (Jensen et al. 2013)

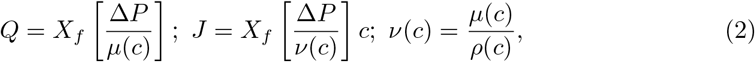

where *X_f_* is a geometric factor that varies with *L* and *a, ΔP* is the pressure difference inside the tube that drives the flow (and need not be only osmotic), *μ*(*c*) is the concentrationdependent dynamic water viscosity that increases exponentially with *c* (Bouchard & Granjean 1995), and *ν*(*c*) is the kinematic viscosity that also increases with *c*. With increasing c, the rise in *μ*(*c*) far exceeds any increase in *ρ*(*c*) so that the functional relations of *μ*(*c*) and *v*(*c*) with *c* are assumed to be the same (within *a* constant *ρ*). For an order of magnitude illustration, increasing *c* from 10% wt/wt to 50% wt/wt increases *ρ*(*c*) by a factor of 1.2 whereas *μ*(*c*) increases by a factor of 4. Because *ν*(*c*) increases non-linearly with *c* as discussed before, a maximum *J* = *J_max_* must exist at a corresponding optimal *c* value that is independent of *X_f_*. Moreover, the existence of this maximum is not predicated based on the precise details of the osmotic controls on Δ*P*. Returning to *J_max_*, for a preset *X_f_*, the hydraulic conductance of the tube *K_t_* can be related to the inverse of viscosity using *K_t_* = *X_f_* / *μ*(*c*). Independent of whether osmotic effects on Δ*P* are fully represented, a *J_max_* associated with an optimal *c* can be derived (numerically here) and graphically shown in figure 1. Moreover, *J_max_* can vary significantly depending on the model presentation within the range of observed values (Jensen et al. 2013).

**Figure 1:**
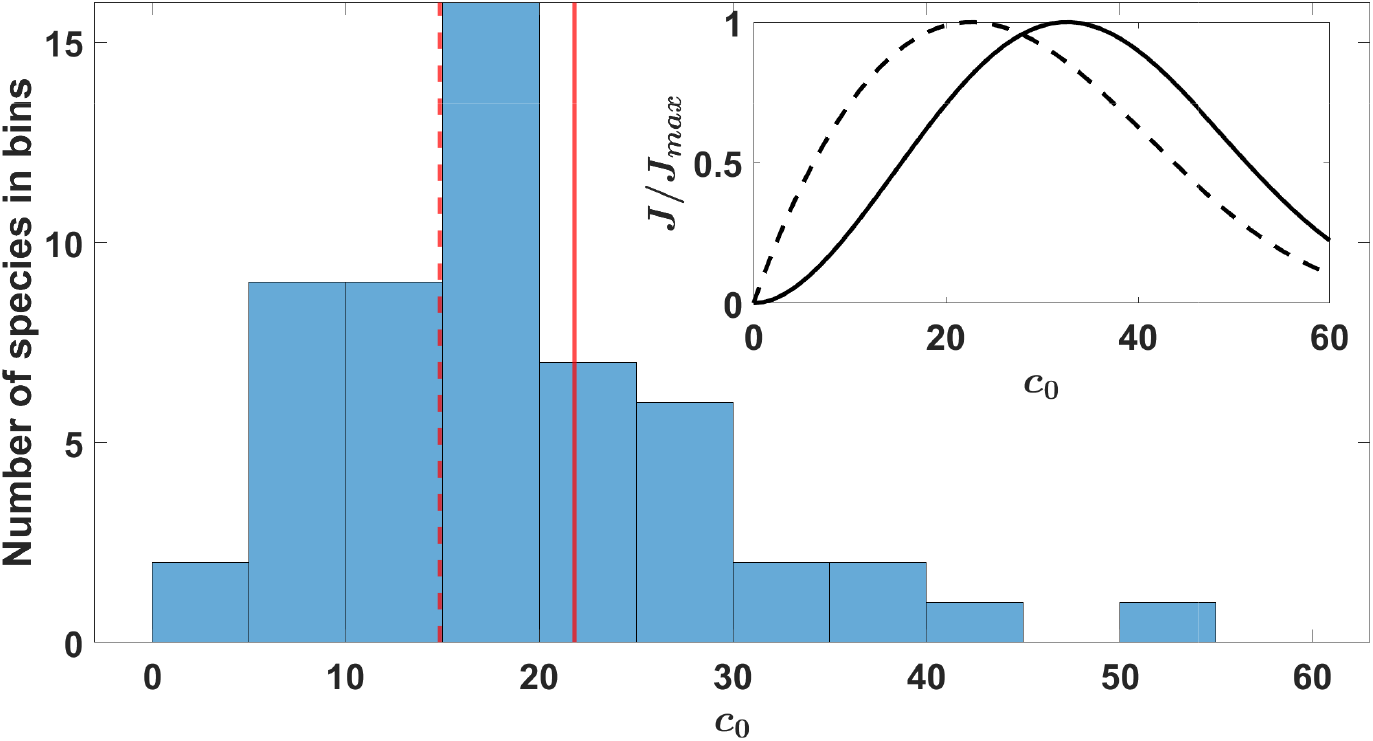
Histogram showing the reported number of species (ordinate) operating at the measured phloem sugar concentration (*c_0_*) values (abscissa) taken from (Jensen et al. 2013). Solid red line (c_0_ ~ 21.8 % wt/wt) denotes the average concentration for active loading species and dashed red line (c_0_ ~ 14.8 % wt/wt) denotes the average concentration for passive loading species. Inset figure shows the computed normalized flux *J/J_max_*for the steady-state and globally averaged Poiseuille models where the solid black line denotes a concentration dependant pressure gradient (through the osmotic relation) and black dashed line denotes an externally imposed constant pressure gradient.

The Van’t Hoff relation approximating the Δ*P* solely from osmotic potential (solid line in figure 1 inset) is given by

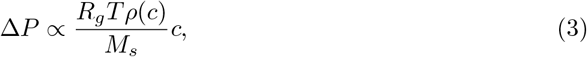

where *R_g_* is the gas constant, *T* is the absolute temperature and *M_s_* is the molar mass of sugar. Similar results with a single *J_max_* but at lower optimum loading sugar concentration (dashed line in figure 1 inset) are produced using an externally supplied constant pressure difference that varies from 1 to 2 MPa with no concentration dependency. This analysis demonstrates that the existence of a *J_max_* is not tightly connected with the Munch hypothesis in the following sense: the precise functional dependence of Δ*P* on *c* is not necessary for the existence of a *J_max_.* Hence, the fact that a *J_max_* exists for a certain *c* is not particularly informative about how axial-radial coordination in the phloem operates and what the role of radial viscosity in this coordination is.

## 2 Results

Transient flow simulations illustrate how local changes, especially near the membrane, lead to further increases in sucrose mass flux above and beyond their steady flow assumption. The choice of a closed end-boundary condition only shows a restrictive case where the osmotic potential decreases because of the spreading of sucrose mass inside the domain. To the contrary, when an open-end boundary condition with infinite sink is used, the sucrose mass flux is expected to be higher because the osmotic potential is constant over the entire simulation. These end-member boundary conditions do not replicate real plants. However, they do amplify the relation between geometry (i.e. long distance transport) and the Munch mechanism as further discussed in section 4.1.1. The results illustrated here show the effect of including *v*(*c*) in the flow equations on mass flux *J*. First, the effect of *v*(*c*) on *J* is discussed by comparing a 2-D model with variable viscosity and a 2-D model with constant viscosity set by the loading concentration (i.e. the two end-member cases discussed in section 4.1). Second, the enhancement of mass transport due to local coordination between axial and radial movement is presented. The results to be featured in this section are generated using numerical simulations for a transient flow with a closed end tube as described in section 4.1.1.

### 2.1 Effect of a concentration-dependent viscosity on *J* (*c*_0_)

This section shows the effect of a concentration-dependent viscosity on sugar mass flux *J* when the loading sugar concentration *c*_0_ is varied. To do so, typical phloem conditions were used to generate the results in both models with constant *v*(*c*_0_) and variable *v*(*c*): *a* = 10*μ*m, membrane permeability *k* = 5 × 10^-14^msPa^-1^ and molecular diffusion coefficient of sucrose in water at standard temperature *D* = 4 × 10^-10^m^2^s^-1^. The *L* was varied from 0.1 to 10 m to describe small or annual plants (e.g. crops) and perennials (trees), respectively. The loading sugar concentration inside the tube *c*_0_ was varied from ~ 3.4 to ~ 61.6 % wt/wt, which does not significantly affect the applicability of the Van’t Hoff relation and the Newtonian fluid approximation for water. For the constant viscosity model, *v*(*c*_0_) does not vary in space and time. The variable viscosity model solves simultaneously for *c* and *v*(*c*) axially and radially in time. The largest signature of viscosity effects on *J/J_max_* is presented in the Supplemental Materials and Methods S3 for results on larger variation in *c*_0_ (up to ~ 83.9 % wt/wt where the viscosity effects may be overestimated due to the over-estimation introduced by the the Van’t Hoff relation and the Newtonian fluid approximation used). In figure 2, the relation between *J/J_max_* and co appears to be similar in shape to the one obtained from the steady and globally averaged Poiseuille model shown in figure 1. However, the optimal concentration at which *J/J_max_* = 1 differs for different tube lengths. Short tube lengths reach *J/J_max_* = 1 at higher *c*_0_ than long tubes. This can be attributed to the role of Taylor dispersion and molecular diffusion in a transient flow simulation not explicitly resolved in a steady and globally averaged Poiseuille model. In this range of *c*_0_, the normalized flux for the variable viscosity model (not shown) exhibits a similar behavior for the *L* = 0.1 m as the constant viscosity case here. However, with increasing L, the optimal point where *J/J_max_* = 1 was not even reached for the range of c_0_ studied (not shown). Interestingly, the normalized flux over a wider range of *c*_0_ for *L* = 2m (selected for illustration) shows that an optimal point does exist for the variable viscosity model as well but c_0_ must operate well outside the range of the Van’t Hoff approximation (see supplemental figure S1B). The fact that *J/J_max_* = 1 was not attained in the variable viscosity model for the range of *c*_0_ selected here may lead to the erroneous conclusion that the inclusion of local viscosity variations in radial and axial directions retards sugar transport. The normalization by *J_max_* hides some facts about the magnitude of *J*, which for the same *c*_0_ is much higher for the variable viscosity model than for the constant viscosity one.

**Figure 2:**
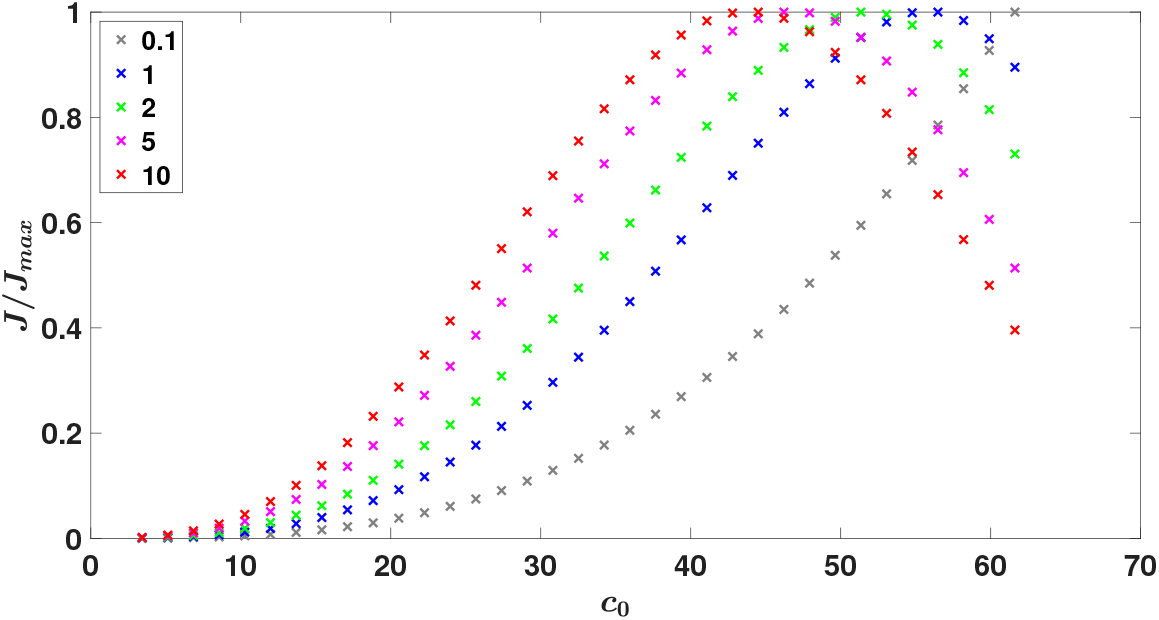
Normalized sugar mass flux (*J/J_max_*) as a function of the initial (or loading) concentration co for the constant viscosity model with *v* = *v*(*c*_0_) in a closed tube. The five-tube lengths (*L* in m) are shown using different colors.

Resolving the local changes in viscosity results in an increase in the overall conductivity of the tube above and beyond the constant viscosity model (figure 3A). The variable viscosity model appears to have far higher *J* than the constant viscosity model at a given *c*_0_ for all tube lengths except for the shortest length (*L* = 0.1m) where the two models are almost indistinguishable. The effect of the tube length is present in both models where an increase in *L* leads to an increase in the flux *J* until a certain value is reached after which *J* starts to decrease with increasing *L*. For example, in the variable viscosity model, the value of *J* when *L* = 10m is lower than the value of *J* when *L* = 5m for the same *c*_0_ as discussed next. An interesting observation is that the value of *L* for which there is a loss of conductivity (the sugar flux decreased for the same initial concentration *c*_0_) is different for both models, *L* = 5m for the variable viscosity model and *L* = 2m for the constant viscosity model.

**Figure 3:**
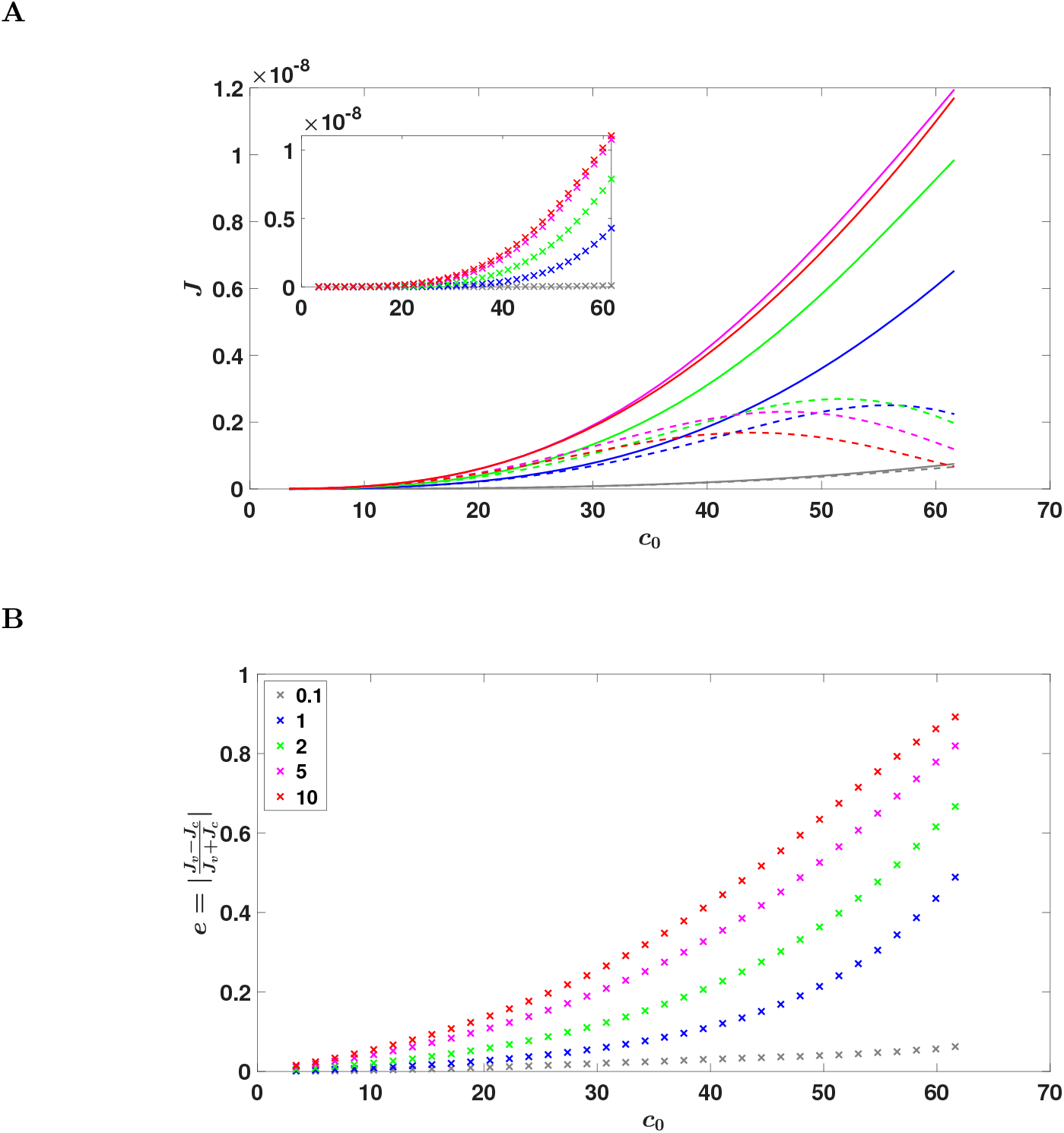
(*a*) Sugar mass flux *J* (Kg s^-1^) as a function of the initial phloem sugar concentration *c*_0_ for the variable viscosity model (solid lines) and the constant viscosity model (dashed lines). Inset plot shows the flux for the viscosity effect (difference between both models). (*b*) Relative difference in sucrose fluxes *e* = | *J_v_ – J_c_|/(J_v_* + *J_c_*) as a function of the initial concentration *c*_0_. Different tube lengths are presented using different colors.

The importance of variable viscosity can be evaluated by the relative difference in sucrose fluxes

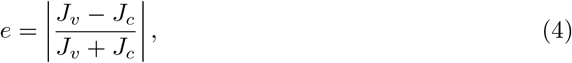

where the subscripts ‘v’ and ‘c’ denote variable viscosity and constant viscosity models, respectively (figure 3B). The results show that *e* increases with *L* and *c*_0_ as expected. This is because the *c*_0_ affects the overall viscosity value itself and *L* affects the development of the velocity profile over which viscosity gradients are allowed to buildup and increase with increasing *L*. The increase in actual mass flux magnitude due to the inclusion of a variable viscosity can be approximated by subtracting the flux of the constant viscosity model from the variable viscosity model (figure 3A inset). As expected, this effect increases with increasing *L*.

### 2.2 Two-dimensional flow results

To understand why the increase in mass flux occurs for the variable viscosity model (vis-a-vis the constant viscosity model), the 2-D simulations are used to illustrate the radial-axial flow and their coordination. These simulations show the local effect of concentration gradients on flow velocity components affected by viscosity and its gradients. Model results show that the computed axial and radial velocities are higher in magnitude because of a lower viscosity near the conduit walls (figures 4A, 4B, 4C). Additionally, the pressure gradient driving the flow is lower compared to the constant viscosity case (figure 4D). The results presented here are for initial concentration *c*_0_ ~ 27.4 % wt/wt and *L* = 2m for the time when the sugar front is located at about 30% of the conduit length, chosen for illustration only. The time *t* it took for the front to reach this location was also different for the two models: *t* = 42.62 s for the variable viscosity model and *t* = 51.05 s for the constant viscosity model (i.e. in the constant viscosity model the flow was about 1.2 times slower). Despite this difference in flow velocity, the sugar front at this location appears similar (4A). The highest difference is near the front location. The velocity profile, on the other hand, appears to be wider, and the difference between the models is higher after the front position (figure 4B). The axial velocity is also higher in the variable viscosity model compared to the constant viscosity model.

**Figure 4:**
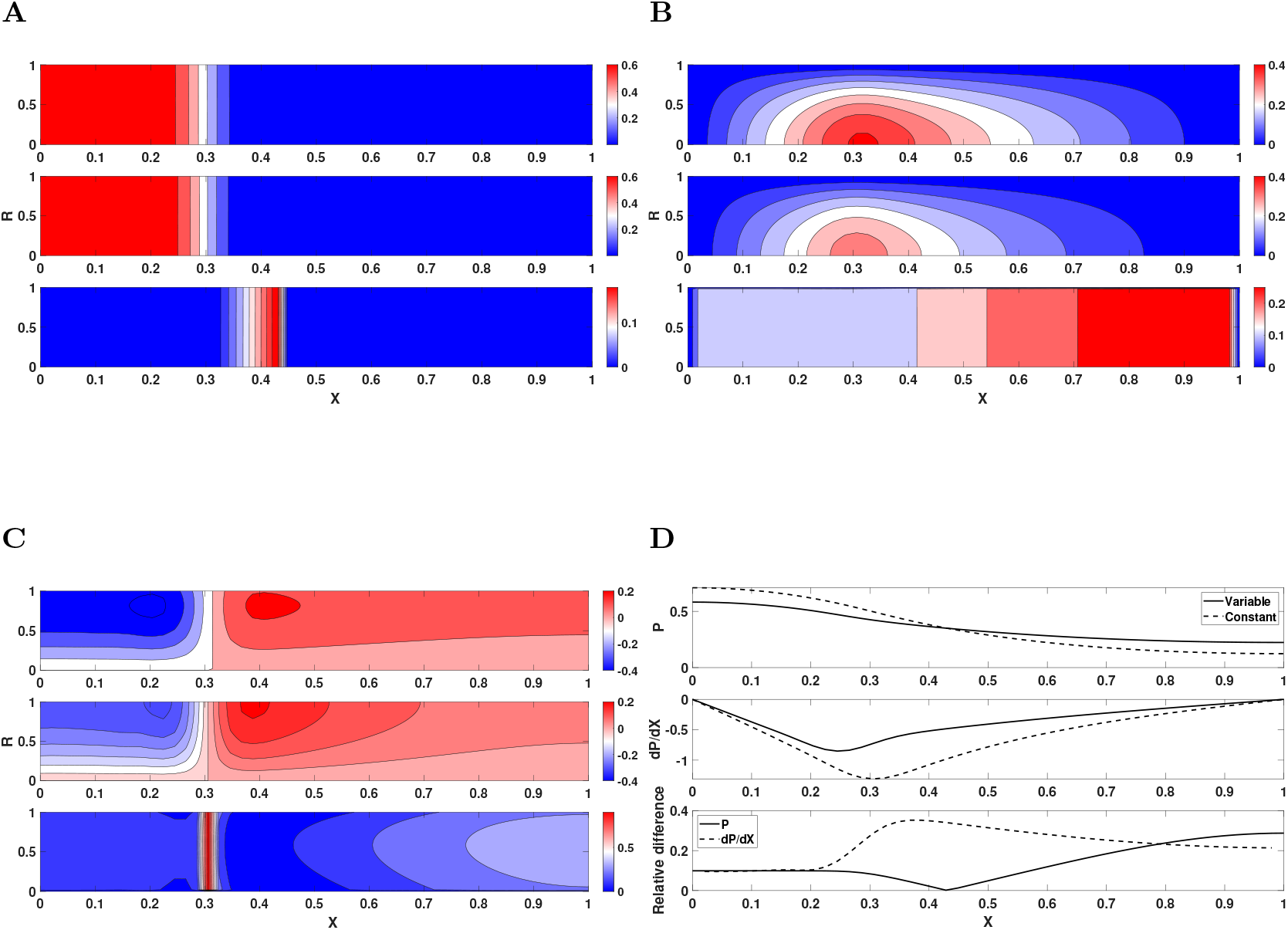
Results for a tube length of *L* = 2m and sugar concentration *c*_0_ ~ 27.4 % wt/wt when the sugar front is located at approximately 30% of the domain. Due to cylindrical symmetry, only half of the domain in the radial direction *R* is shown. (*a*) concentration profiles, (*b*) axial velocity profiles, and (*c*) radial velocity profiles for the variable viscosity model (top), constant viscosity model (middle), and the relative difference (bottom), respectively. (*d*) pressure (top figure), pressure gradient (middle figure), and their relative difference (bottom figure) for the variable viscosity and constant viscosity models. The relative difference between the models is calculated by *e*.

The effect of local viscosity gradients generated by concentration gradients is more apparent in the radial velocity than the axial velocity as expected (figure 4C). This finding can be anticipated from equation (11) since the radial velocity profile is directly related to viscosity gradients in the axial direction. These gradients are generated based on concentration gradients from the axial direction that are large, due to the wave nature of the problem (because the advection-diffusion equation has a wave shape especially when it is advection-dominated). The relative difference e between the models is high near the front location. Moreover, the location of the maximum radial velocity is shifted further away from the membrane for the variable viscosity model since the tube conductance has a new term that depends on non-local changes in the radial direction (i.e. (*K_t_*)*_r_* in equation (10)). Similar to the axial velocity profile, this non-local effect is also apparent in the radial velocity profile for the variable viscosity model that has a wider velocity range, when compared to the constant viscosity model. Due to a lower sugar concentration near the membrane, the viscosity of the sap decreases leading to less resistance to the radial inflow of water (that is the driving force for osmotically driven flows) in the variable viscosity model. This can be conceptually understood as a decrease in wall-friction when area-averaging the equations.

The variable viscosity model also has a smoother pressure field compared to the constant viscosity model (figure 4D). An interesting result in this figure is that the global pressure gradient over the domain is smaller for the variable viscosity model and yet the front travels at a higher speed. This paradoxical result can only be explained by the increase in conductivity of the tube because of local viscosity effects being coordinated in radial and axial directions.

## 3 Discussion

In this work, the Münch mechanism is revised to include viscosity variations due to concentration variations in the axial and radial directions within the phloem. Numerical simulations were conducted to assess the effects of these additions over the conventional approach of setting a single viscosity value that depends on the loading concentration. These simulations show that the inclusion of viscosity variations with concentration leads to a speed-up in the phloem sap flow when compared to a constant viscosity model. The variable viscosity model suggests more resiliency may be achieved when compared to the constant viscosity model because higher flow rates can be achieved with smaller overall pressure gradients. Furthermore, plant optimal *L* is expected to drop faster with increasing loading concentration for the constant viscosity model than for the variable viscosity model. This result is apparent from the data shown in figure 1 where tall trees don’t have as high sugar concentrations in the leaves as would be expected for plants with long phloem transport pathways.

### 3.1 Model refinement leads to a speed up in mass flux

The concentration-dependent viscosity has two effects on *J* - both leading to an increase in its magnitude. The first is a smaller viscosity in the vicinity of the membrane since the water influx dilutes the solution near the membrane. This process is also accompanied by an opposite process where the decrease in c near the membrane leads to a lower osmotic effect and radial velocity (Pedley 1983). However, in the simulations here, the osmotic forcing is dynamic and dictates the concentration at the membrane through a dynamic boundary condition. This dynamic boundary condition is formulated using Darcy’s law discussed in section 4.1.1 where the effect of this ‘unstirred’ layer is accounted for. This unstirred layer is further discussed in the Supplemental Materials and Methods S4 where the thickness of this layer was estimated using the method discussed in Pohl et al. (1998) and is reasonably resolved in the simulations. The total effect of the aforementioned processes considered here remains an increase in *v* near the membrane. This increase has the effect of radially advecting sugar molecules away from the pipe walls and towards the pipe center. Because the axial velocity profile peaks in the pipe center and yet the radial viscous stress is zero there, the overall mass flux is also increased. The second is that the increase in radial velocity near the pipe walls, when coupled to a zero radial velocity at the pipe center (as required by symmetry), must be accompanied by an increase in axial velocity gradients (due to the incompressibility approximation). Thus, a speeding up of *u* is expected. The analysis of the axial pressure distribution further suggests that this effect is sensed over a broader region of *L*. This speeding up yields a faster front advancement of sugar away from the loading zone for the variable viscosity model. Both mechanisms are operating in concert to increase J above and beyond the constant viscosity model. That those two effects act together to enhance tube permeability affecting *J*, not the driving force for water (i.e. pressure gradients) is also supported by the analysis here. For this reason, the *J_max_* analysis and its dependence on *c*_0_ leading to *c*_0_ ≈ 20% wt/wt in figure 1 being not sensitive to *X_f_* (as earlier shown) cannot detect this coordination between axial and radial flow.

### 3.2 Implications to validity of the MUnch mechanism in plants

For the purposes of representing sucrose inflow into the phloem, plants can be naively divided into two categories: small *L* plants such as herbaceous plants or crops that are mainly active sugar loaders, resulting in more efficient use of photoassimilates, and large *L* plants such as trees exhibit the characteristics of passive sugar loaders. Figure 1 shows that active loaders typically have higher loading concentration (*c*_0_ ~ 22 % wt/wt) than passive loaders (*c*_0_ ~ 15 % wt/wt) for all the species considered in each category. At first glance, this observation seems to contradict the basic assumption of Münch mechanism, because high c0 increases viscosity and can potentially slow down phloem transport. Pressure gradients driving water (i.e. HP equation) and sugars down the phloem, when formulated as pressure differences between sources and sinks normalized by *L* must require higher *c_0_* to compensate for the larger *L* to attain the same flow velocity. The higher *c_0_* increases the driving pressure at the loading site linearly (when given by the Van’t Hoff relation). However, this argument is simple at best. From the supplemental figure S2, the maximum *J* = *J_max_* exists at an optimal length tube *L* value that decreases with increasing loading concentration *c_0_* for both models. For the constant viscosity model, the optimal *L* drops faster than the variable viscosity model. The results from the analysis here offers a new perspective on this issue. Consider an active loader such as maize that has typical *c*_0_ ~ 50 % wt/wt and is 3-4 m tall (= L). Using this *c_0_* and inspecting figure 3A, the variable viscosity model predicts a maximum *J* at *L*=5 m whereas the constant viscosity model predicts a maximum *J* for an *L* = 2 m only. Now when passive loaders are considered, the variable viscosity model shows more resilience for increasing phloem length when compared to the constant viscosity model. This can be seen from figure 3B where for low loading concentration, increasing the length of the tube leads to a higher relative difference between the models, meaning the sucrose mass flux is higher for the variable viscosity model compared to the constant viscosity model. These findings imply that using simplified physics with a constant viscosity underestimates the speed of the flow leading to erroneous conclusions on long distance transport. In addition, the effect of a concentration-dependent viscosity on sucrose transport in phloem resulted in a lower pressure gradient driving the flow along the axial direction. In active phloem loaders, the price to pay in terms of restricted height or path length maybe outweighed by the benefits of rapid growth and the possibility of competing with each other. This finding contributes to the growing evidence that the pressure-flow hypothesis can provide the necessary mechanism for long-distance sugar transport as long as the complexity in transport physics is accommodated. The work here also showed that viscosity adjustments lead to conductivity enhancement for sugar transport instead of pressure gradients for water flow. This is shown in figure 4D where for the same loading sugar concentration, a smaller pressure gradient along the phloem pathway is generated for the variable viscosity model, yet a higher sap speed because of the increased conductivity. These findings may offer new hypotheses about the resilience of plants during drought condition. More negative xylem water tension requires a higher osmotic potential to maintain phloem transport in dry conditions (Sevanto 2014). In this case where a high osmotic potential is needed, the variable viscosity model shows that the phloem can still function under these conditions without failure because the pressure gradient generated to drive the flow was lower than the one predicted from the constant viscosity model. In addition, the increase in optimal sugar concentration operating range for *J* would allow plants to increase their sucrose concentration to potentially overcome those large tensions arising in the xylem during drought without a substantial decrease in mass flow rate. Future work will focus on the effects of nonlinear xylem water potential on phloem transport by including sugar sources and sinks along the phloem path.

## 4 Materials and methods

To isolate the effect of a concentration-dependent viscosity on radial-axial flow coordination, many simplifications must still be invoked when representing the physics of translocation in a cylindrical tube. In all formulations considered here, it is assumed that i) the phloem vasculature can be approximated by a long slender tube of length *L* and radius *a* (*∈* = *a*/*L* << 1) with rigid semipermeable walls characterized by a constant permeability *k* that allows the exchange of water molecules but not sugars with the surroundings, ii) sieve plates have minimal effect on the flow and can be modeled as either an ‘extra’ drag force uniformly acting along with L or ignored altogether, (iii) the bulk flow is at very low Reynolds number *Re* ~ *pauμ*^-1^ ≪ 1 where u is a characteristic longitudinal velocity, *μ* the dynamic viscosity, and *ρ* the density, so that creeping flow is maintained throughout, iv) sugar sources and sinks are modeled as boundary conditions at the entry and exit end of the tube. Hence, water can be exchanged with the surroundings but not sugars thereby suppressing any enhancement due to a relay effect.

### 4.1 Variable Viscosity Model

To derive the general model that includes concentration and viscosity variations in axial and radial directions, the governing equations under certain assumptions and simplifications are first analyzed in Cartesian coordinates. Then, the non-dimensional form of these equations, which are necessary for analyzing the numerical model results, are presented in cylindrical coordinates. The model to be presented in this section is considered one ‘end-member’ case in including a concentration-dependent viscosity. The other ‘end-member’ case this model is compared to is the constant viscosity model that assumes a single viscosity value in the domain set by the initial loading concentration. An example of this model for globally averaged steady-state conditions is discussed by Jensen et al. (2013). Thompson & Holbrook (2003) present a different model that is between these ‘end-member’ cases. In their work, they included local variations in viscosity inside the domain but only using radially averaged equations (i.e. variations in viscosity along radial directions ignored). Due to the nonlinear relation in the viscous stress between velocity and viscosity, area-averaging the equations leads to a simplified model that excludes the effect of this non-linearity. This issue can be resolved at the expense of solving the equations in axial and radial directions and frames the main approach here.

#### 4.1.1 Governing equations

In a three-dimensional Cartesian coordinate system defined by longitudinal (*x_1_* = *x*), lateral (*x*_2_ = *y*), and vertical (*x_3_* = *z*) directions, water flow within the phloem satisfies the continuity equation

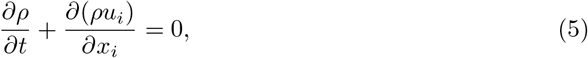

where *t* is time, *i* = 1, 2, 3 describe direction *x_i_*, and *u_i_* is the instantaneous velocity along direction *x_i_*. Here, index notation is used with repeated indices implying summation unless otherwise stated. The flow must also satisfy the conservation of momentum, which describes the force balance along direction *x_i_*, and is given as

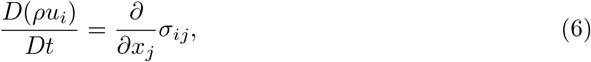

where *D*(.)/*Dt* is the material derivative and *σ_ij_* are the nine components of the stress tensor. The *σ_ij_* of a Newtonian fluid can be approximated by

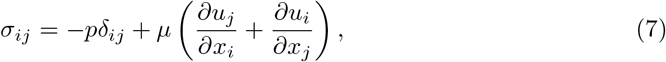

where *p* and *μ* are the local pressure and dynamic viscosity of the fluid, respectively, and the *δ_ij_* is the Kronecker delta (i.e. *δ_ij_* = 0 when *i* ‡ *j* but unity otherwise). This representation of *σ_ij_* is approximate and assumes that the stress tensor is symmetric (*σ_ij_* = *σ_ji_*) and the so-called second viscosity coefficient (or volume viscosity) is momentarily ignored (Panton 2006). Equation (7) also assumes that there are no external forces on the fluid and that the gravitational forces cancel out (Thompson & Holbrook 2003).

In terms of *fluid properties*, *ρ* depends on c and strictly speaking cannot be treated as constant when *c* varies in time or along *x_i_*. However, this dependence is minor when compared to variations in *μ* as demonstrated earlier (see section 1) and variations in *ρ* will be assumed small for simplicity so that *∂u_i_/∂x_i_* = 0. In this case, the *σ_ij_* representation given by equation (7) is reasonable (Panton 2006). Another common assumption in phloem transport is that *μ* is constant set by the loading concentration. This approximation is only applicable for small concentration values. However, in plants, *c* can range from 15% wt/wt to 35% wt/wt (and for maple trees even up to ≈ 50% wt/wt). In this high concentration range, the dependence of viscosity on concentration has not been fully analyzed in the context of three-dimensional water and sugar transport. Some models include this dependence of *μ* on *c* in an area-averaged formulation (Thompson & Holbrook 2003, Jensen et al. 2016), but area-averaged formulations that evolve concentration axially and presume uniform concentration along the radial direction cannot resolve radial-axial flow coordination to be studied here. Therefore, the model proposed here includes the dependence of *μ* on *c* in both axial and radial directions and tracks its consequences on the shape of the *J*–*c* relation as well as the magnitude of *J* across differing *L* and loading concentrations.

In terms of *flow properties*, the low Reynolds number (Re ≪ 1) and small aspect ratio (e ≈ 10^-5^) can be used to show that under the so-called lubrication theory (where the flow in one of the dimensions is significantly smaller than the others because of geometric constraints), the momentum balance may be approximated by (now written in cylindrical coordinates)

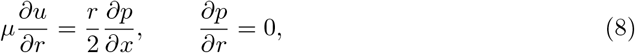

where *x* remains the axial direction with *x* = 0 situated at the loading zone, *r* is the radial direction with *r* = 0 defining the center of the tube, and *u*(*x*, *r*) and *v*(*x*, *r*) are the axial and radial velocity components respectively at any point (*x*,*r*). In Supplemental Materials and Methods S1, the derivation of this formulation and all its assumptions are presented for completeness.

To understand how viscosity gradients affect the flow, equation (8) can be written in a compact form. First, the tube conductance is re-defined as *K_t_* = 1/μ as in section 1 (the geometric factor *X_f_* is absent since it is the result of radial averaging). Integrating equation (8) in the radial direction while noting that the pressure *p* is only a function of the axial position (as shown in Supplemental Materials and Methods S1) leads to

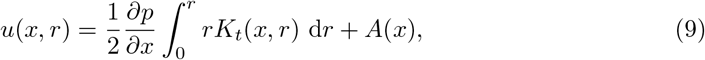

where *A*(*x*) is an integration function that varies in *x*. Using the no-slip boundary condition on the longitudinal velocity component *u*(*x*, *a*) = 0 at the membrane surface set as *r* = *a* leads to *A*(*x*) = – (*a*^2^〈K_t_〉/4) (*∂p/∂x*) where 〈*K_t_*〉 is the radially averaged tube conductance. Similarly, the term 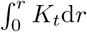 can be written as *r^2^*〈*K_t_*〉*_r_*/2 where the subscript *r* denotes radial averaging until the current radial position (*r* ≤ *a*). Using this form, equation (9) describes the axial velocity

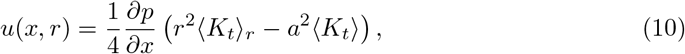

where both terms 〈*K_t_*〉*_r_* and 〈*K_t_*〉 are functions of *x* but only 〈*K_t_*〉*_r_* is a function of *r*.

Equation (10) is different from the HP expression because the variable viscosity depends on *c*(*x*, *r*) that itself varies radially and axially. To be clear, this dependence is nonlocal because of the integral operator in the r direction. However, if a constant viscosity is used at a given *x*, 〈*K_t_*〉*_r_* and 〈*K_t_*〉 are equal to 1/*μ*, and the aforementioned conservation of momentum equation becomes equivalent to the HP expression with an adjustment. This adjustment is due to osmosis that generates a radial inflow of water leading to *∂*^2^*p*/*∂x*^2^ = 0, which then leads to a variable pressure gradient instead of a constant one as is common in HP applications in pipes (Phillips & Dungan 1993). However, the partial *∂p*/*∂x* not being constant does not violate or invalidate the HP equation as discussed elsewhere (Thompson & Holbrook 2003, Nakad et al. 2021).

The effect of viscosity gradients is not directly apparent in equation (10). To make it explicit, the continuity equation (5) in cylindrical coordinates is now considered. It is given as

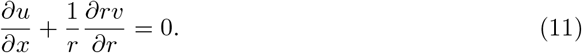

Here *ρ* variation with *c* is once again assumed to be small compared to the viscosity variations with *c* as stated before. Using the expression for the axial velocity from equation (10) in the continuity equation (11), one can see how axial viscosity gradients impact the radial velocity *v*, which is not identically zero due to osmosis. Moreover, the viscosity gradient is not only the result of the area-averaged tube conductance 〈*K_t_*〉 but also stems from the radially-averaged (or non-local) tube conductance 〈*K_t_*〉*_r_* that depends on radial position *r*. Equation (8) with equation (11) can be used to describe the flow of water characterized by *u*(*x*,*r*) (axial velocity) and *v*(*x*,*r*) (radial velocity) inside the tube as a function of position *x*, *r*.

Equations (8) and (11), however, remain incomplete since there are two equations with three unknowns *u*, *v*, and *p*. This mathematical setup is in sharp contrast to flow in closed pipes where *v* =0 everywhere due to the solid wall boundary condition at *r* = *a* and symmetry considerations at the pipe center. In phloem, osmosis necessitates a finite *v* at the pipe walls while symmetry considerations alone result in *v* = 0 at the center of the pipe. Thus, the third equation that relates *v* to total pressure inside the tube must be provided by osmoregulation. This equation is best formulated as a boundary condition describing a flow through a porous media (a thin membrane here) at *r* = *a*. Such a boundary condition may be given by a Darcy-type formulation assuming a very low Reynolds number for the radial flow into or out of the pipe walls. This boundary condition yields an expression for *v* at *r* = *a* given by

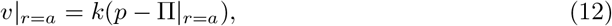

where Π|*_r_*=*_α_* is the osmotic potential at the membrane and *k* is the membrane permeability assumed constant and independent of *v* (i.e. no Forscheimer or quadratic corrections to Darcy’s law). This osmotic potential can be related to *c* using the Van’t Hoff relation, Π = *R_g_Tc*(*x*, *a*) as before. This approximation is reasonable for low c and compatible with the assumption that the density and molecular diffusion (discussed below) do not vary appreciably with *c* when compared to viscosity.

The last equation needed to describe the physics of sugar transport is the conservation of solute mass, which is also needed to solve for *u*, *v*, and *p*. This equation is derived using Reynolds transport theorem that describes the movement of solutes (mainly sugar here) due to advection and molecular diffusion. In cylindrical coordinates, it is given by

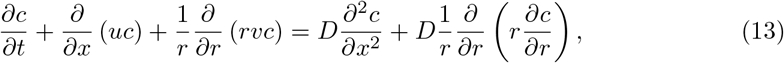

where *D* = *vSc*^-1^ is the molecular diffusion coefficient of sugar in water assumed to be again insensitive to *c* variations when compared to *ν*, and *Sc* ≫ 1 is the molecular Schmidt number for sugars in water (usually of order 10^4^).

The final step for describing the physics of sugar transport is to specify the boundary and initial conditions. These are problem-specific and are selected here to illustrate one restrictive ‘end-member’ case of flow in a closed tube with no sugar sinks. This case is dynamically interesting because sugar concentration keeps building up as no sugars are removed from the tube. The other ‘end-member’ case is where sugars are instantly consumed at the end of the pipe (i.e *c*(*L*,*r*) = 0 and sugar sinks are treated as ‘infinite’). This latter case is expected to lead to a much larger J in the tube, which is why the focus is on the more restrictive former case. In plants, *c*(*L*)/*c*(0) ≪ 1 and thus osmotic gradients are much higher in the presence of sinks than those set by the closed tube assumption. Thus, the physics of closed tubes must require higher loading concentrations to drive the water velocity, which is why they are more restrictive and thus dynamically interesting from the perspective of exploring limitations on the Munch hypothesis. In a pipe closed at both ends *u*(*x* = 0) = *u*(*x* = *L*) = 0, water flow must accelerate to a well-defined maximum and then decelerate to zero velocity along *L*. For initial conditions selected here, sugar is released as an axially smooth function *c*(*x*, *t* = 0) = *f*(*x*) with no radial variation, meaning that radial diffusion is initially fast enough to ensure a uniform distribution of sucrose along *r*. The closed tube assumption with no sinks requires sugar mass to be conserved inside the tube during the entire period resulting in

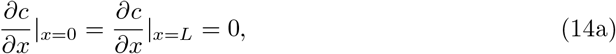

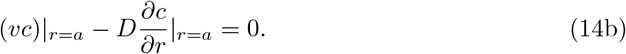

Finally, a no-slip boundary condition at the membrane in the axial direction only, i.e. *u*|_*r*=*a*_ = 0, and symmetry considerations at the center of the tube, where *v*|_*r*=0_ = 0 and *∂c*/*∂r*|_*r*=0_ = 0, are all enforced.

#### 4.1.2 Non-dimensional form and key dimensionless numbers

To elucidate the key dimensionless numbers governing water and sugar movement, and make interpretation of the equations easier, this section describes the scaling analyses and the non-dimensional form of the equations used. To write the equations in dimensionless form, the following scales were adopted: The *x* and *r* were scaled by the tube length L and radius *a*, respectively, leading to *x* = *LX*, *r* = *aR*. Time, and axial and radial velocity as well as pressure and concentration were scaled by their respective initial values at *x* = 0 (subscript 0) leading to *t* = *t_0_τ*, *u* = *u_0_U*, *v* = *v_0_V*, *p* = *p_0_P*, *c* = *c_0_C* and 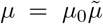. The dynamic viscosity *μ* was scaled by *μ*_0_, determined from *c*_0_ and *T*. This *μ*_0_ will also be used in a constant viscosity model as a reference to assess the impact of accommodating variable viscosity in *σ_ij_* and its spatial gradients. Using these scales, the non-dimensional equations for the two velocity components and pressure are

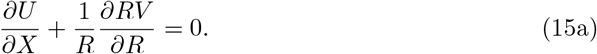

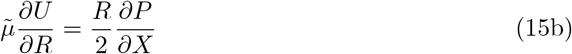

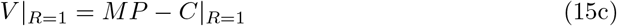

where *u*_0_ = *kR_g_*Tc_0_∈^-1^, *v_0_* = *∈u*_0_, *p*_0_ = *μ*_0_*Lu*_0_*a*^-2^, and *M* = *kμ*_0_*L*^2^*a*^-3^ is the Munch number defined as the ratio of axial resistance over radial resistance as discussed in (Jensen et al. 2009, Nakad et al. 2021). The complete scaling analysis for the Navier-Stokes equations is shown in Supplemental Materials and Methods S1.

The non-dimensional form of the conservation of sugar mass, i.e. equation (13), is

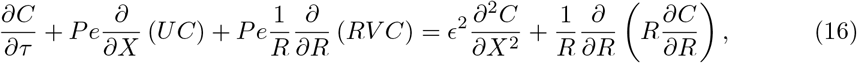

where *t*_0_ = *a*^2^*D*^-1^ is the radial diffusion timescale and *Pe* = *v*_0_*aD*^-1^ is the radial Peclet number defined by the ratio of radial advection to radial diffusion. This expression explicitly shows the relative contributions of radial flow dynamics (through *Pe*) and simplified geometry (through the slender ratio *∈*) to mass transport.

### 4.2 Model calculations

The proposed model calculations provide the variations of *J*(*x*, *r*), *c*(*x*,*r*), *u*(*x*, *r*), *v*(*x*, *r*), and *p*(*x*, *r*) at every *t*. A description highlighting the numerical method used to obtain the results is presented in Supplemental Materials and Methods S2. To link a representative *J* with a *c*_0_ in a manner that allows comparison with the prior *J_max_* and optimal *c* analysis, the following steps and approximations were taken in post-processing the model results. With a radial Peclet number *Pe* ≪ 1, it is reasonable to assume that the mass flow primarily occurs in the axial direction (Nakad et al. 2021). The molecular diffusion can also be neglected since the axial Peclet number, defined by the ratio of axial advection to axial diffusion and derived from *Pe* by *Pe_l_* = *∈*^-2^*Pe*, is large (i.e. *Pe_l_* ≫ 1). Using these assumptions, the area-averaged sugar flux can be reasonably de termined from post-processing time variations of the sugar front position *x_f_*. This front is also delineated from maximal |*∂c*(*x*, *r*)/*∂x*|. To determine *x_f_*, we fitted an exponential relation between *x_f_* and *t* using (Jensen et al. 2009, Nakad et al. 2021)

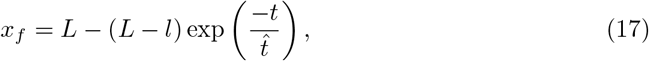

where *l* ≈ 0.2 is the initial sucrose front position, *L* = 1 is the length of the tube and *x_f_* is the front position all in non-dimensional form. Subtracting *l* from both sides of equation (17), a linear relation between *t* and ln [(*X_final_* – *X_f_*) / *X_final_*] (where *X_fina_* = *L* – *l* ≈ 0.8 and *X_f_* = *x_f_* – *l*) can be obtained. Hence, linear regression applied to the 2D numerical solution was then used to obtain the constant 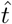 away from the entrance boundary condition. The front speed was determined as *U_s_* = *dx_f_* / *dt*. Finally, the mass flux was approximated by *J_num_* = *Q_num_C* where *Q_num_* is the numerical volumetric flow rate. The *Q_num_* was determined using approximated speed *Q_num_* = *A_t_U_s_* where *A_t_* is the cross-sectional area of the tube. We present results from the two-dimensional (2-D) model simulation for the axial velocity *U*, radial velocity *V*, concentration *C*, and pressure *P* in the dimensionless form to illustrate the effect of variable viscosity on radial and axial variations of these variables. The model simulations were conducted using MATLAB programming language (Mathworks, Natick, MA).

## Acknowledgments

MN, J-CD, and GK acknowledge support from the U.S. National Science Foundation (NSF-IOS-1754893, NSF-AGS-2028633) and the Department of Energy (DE-SC0022072). SS was supported by Los Alamos Directed Research and Development Exploratory Research Grant (No. 20160373ER). No data are generated or used in this work.

