## Supplemental Materials and Methods for "Radial-axial transport coordination enhances sugar translocation in the phloem vasculature of plants"

### Materials and Methods S1 Model Derivation

In section 4.1.1, the variable viscosity model was derived using the lubrication theory approximation. Its salient features are provided here. Starting from the assumption that sap flow can be described as an incompressible Newtonian fluid and using the relation in equation (7) for the stress tensor, the conservation of momentum in equation (6) can be written as

$$\rho \frac{Du_i}{Dt} = -\nabla_i p + \nabla \cdot \left[\mu \left(\nabla_j u_i + \nabla_i u_j\right)\right], \tag{S.1}$$

or in expanded form

$$\rho \frac{Du_i}{Dt} = -\nabla_i p + \mu \nabla \cdot \left(\nabla_j u_i + \nabla_i u_j\right) + \nabla \mu \cdot \left(\nabla_j u_i + \nabla_i u_j\right), \tag{S.2}$$

which leads to

$$\rho \frac{Du_i}{Dt} = -\nabla_i p + \mu \nabla \cdot \nabla u_i + \mu \nabla_i \nabla_j \cdot u_j + \nabla_j \mu \nabla_j u_i + \nabla_j \mu \nabla_i u_j. \tag{S.3}$$

Equation (S.3) can be further simplified by setting the third term to zero due to the incompressibility assumption. Noting that the second term on the right-hand-side is the Laplace of $u_i$ ($\nabla \cdot \nabla = \nabla^2$), the conservation of momentum reduces to

$$\rho \frac{Du_i}{Dt} = -\nabla_i p + \mu \nabla^2 u_i + \nabla_j \mu \nabla_j u_i + \nabla_j \mu \nabla_i u_j. \tag{S.4}$$

Assuming cylindrical symmetry, the momentum balance in the axial and radial directions can be written, respectively, as

$$\begin{aligned}
\rho\left(\frac{\partial u}{\partial t} + u\frac{\partial u}{\partial x} + v\frac{\partial u}{\partial r}\right) &= -\frac{\partial p}{\partial x} + \mu\left[\frac{\partial^2 u}{\partial x^2} + \frac{1}{r}\frac{\partial}{\partial r}\left(r\frac{\partial u}{\partial r}\right)\right] \\
&\quad + 2\frac{\partial \mu}{\partial x}\frac{\partial u}{\partial x} + \frac{\partial \mu}{\partial r}\frac{\partial u}{\partial r} + \frac{\partial \mu}{\partial r}\frac{\partial v}{\partial x}, \\
\rho\left(\frac{\partial v}{\partial t} + u\frac{\partial v}{\partial x} + v\frac{\partial v}{\partial r}\right) &= -\frac{\partial p}{\partial r} + \mu\left[\frac{\partial^2 v}{\partial x^2} + \frac{1}{r}\frac{\partial}{\partial r}\left(r\frac{\partial v}{\partial r}\right) - \frac{v}{r^2}\right] \\
&\quad + 2\frac{\partial \mu}{\partial r}\frac{\partial v}{\partial r} + \frac{\partial \mu}{\partial x}\frac{\partial v}{\partial x} + \frac{\partial \mu}{\partial x}\frac{\partial u}{\partial r}.
\end{aligned} \tag{S.5}$$

The continuity equation can also be written in cylindrical coordinates as

$$\frac{1}{r}\frac{\partial (rv)}{\partial r} + \frac{\partial u}{\partial x} = 0. \tag{S.6}$$

The scaling variables defined by $u = u_0 U$, $v = v_0 V$, $p = p_0 P$, $r = aR$, $x = LX$ and $\mu = \mu_0 \tilde{\mu}$, where $u_0$, $v_0$ and $p_0$ are characteristic axial velocity, radial velocity and pressure respectively. The characteristic length scales are the length and radius of the tube $L$ and $a$, respectively. The characteristic viscosity scale $\mu_0$ is related to the initial concentration $c_0$ (i.e. the loading concentration). The radial velocity scale $v_0 = \epsilon u_0$ is determined from the continuity equation, and $p_0 = (L\mu_0 u_0)/a^2$ is the viscous pressure scale. Using these normalized variables, the non-dimensional form of the continuity and Navier-Stokes equations, while assuming steady-state creeping flow, are

$$\begin{aligned}\frac{1}{R}\frac{\partial(RV)}{\partial R}+\frac{\partial U}{\partial X}&=0,\\ \epsilon Re\left(U\frac{\partial U}{\partial X}+V\frac{\partial U}{\partial R}\right)&=-\frac{\partial P}{\partial X}+\tilde{\mu}\left[\epsilon^2\frac{\partial^2 U}{\partial X^2}+\frac{1}{R}\frac{\partial}{\partial R}\left(R\frac{\partial U}{\partial R}\right)\right]\\ &\quad+2\epsilon^2\frac{\partial\tilde{\mu}}{\partial X}\frac{\partial U}{\partial X}+\frac{\partial\tilde{\mu}}{\partial R}\frac{\partial U}{\partial R}+\epsilon^2\frac{\partial\tilde{\mu}}{\partial R}\frac{\partial V}{\partial X},\\ \epsilon^3 Re\left(U\frac{\partial V}{\partial X}+V\frac{\partial V}{\partial R}\right)&=-\frac{\partial P}{\partial R}+\epsilon^2\tilde{\mu}\left[\epsilon^2\frac{\partial^2 V}{\partial X^2}+\frac{1}{R}\frac{\partial}{\partial R}\left(R\frac{\partial V}{\partial R}\right)-\frac{V}{R^2}\right]\\ &\quad+2\epsilon^2\frac{\partial\tilde{\mu}}{\partial R}\frac{\partial V}{\partial R}+\epsilon^4\frac{\partial\tilde{\mu}}{\partial X}\frac{\partial V}{\partial X}+\epsilon^2\frac{\partial\tilde{\mu}}{\partial X}\frac{\partial U}{\partial R},\end{aligned}\tag{S.7}$$

where $Re=\rho u_0 a\mu_0^{-1}$ is the Reynolds number. Under the lubrication theory approximation, the leading order equations become

$$\tilde{\mu}\frac{1}{R}\frac{\partial}{\partial R}\left(R\frac{\partial U}{\partial R}\right)+\frac{\partial\tilde{\mu}}{\partial R}\frac{\partial U}{\partial R}=\frac{\partial P}{\partial X},\qquad \frac{\partial P}{\partial R}=0.\tag{S.8}$$

The left-hand side of equation (S.8) can be further simplified by multiplying the equation by $R$ leading to

$$\tilde{\mu}\frac{\partial}{\partial R}\left(R\frac{\partial U}{\partial R}\right)+\frac{\partial\tilde{\mu}}{\partial R}R\frac{\partial U}{\partial R}=R\frac{\partial P}{\partial X},\tag{S.9}$$

or written in standard form,

$$\frac{\partial}{\partial R}\left(\tilde{\mu}R\frac{\partial U}{\partial R}\right)=R\frac{\partial P}{\partial X}.\tag{S.10}$$

Equation (S.10) can now be integrated in the radial direction and leads to

$$\tilde{\mu}\frac{\partial U}{\partial R}=\frac{R}{2}\frac{\partial P}{\partial X},\tag{S.11}$$

where the constant of integration was set to zero due to symmetry consideration (i.e. $\partial U/\partial R = 0$).

### Materials and Methods S2 Numerical Scheme

A brief description of the numerical method used to obtain the results discussed in section 2 is presented. Spatially, the tube is divided into equally spaced grids where the number of grids in axial and radial directions was set to 50 in each direction. The calculations were checked to ensure the solution is independent of the grid size by doubling the grid points for the large $M$ (or long tube). Initially, the concentration profile in the tube was assumed to be uniform along with $r$ but maintains a specified front shape along the $x$-direction through the following relation:

$$C(X,R,\tau=0)=f(X)=1-\left[1+\exp\left(\frac{\lambda-X}{\alpha}\right)\right]^{-1},\tag{S.12}$$

where $\lambda$ and $\alpha$ are two defined constants that describe the initial size and smoothness of the function. Here, $\lambda$ and $\alpha$ were chosen to be 0.2 and 0.02, respectively. From this initial condition, the dynamic viscosity at $\tau = 0$ can be solved using standard equations (Bouchard & Granjean 1995). This allowed the velocity and pressure field to be computed using equations (15a), (15b), and (15c) where the boundary condition $\partial(RV)/\partial R = 0$ at $R = 1$ was imposed. A central finite difference method was used for spatial derivatives. After solving for the initial velocity and pressure, an implicit scheme using Newton's method was then used to integrate equation (16) in time.

### Materials and Methods S3 Extrapolating to Extreme Sugar Concentration

In this section, the effect of high initial concentration on the models is discussed. For high sugar concentrations, the Newtonian fluid and Van't Hopf assumptions can be questionable. However, to extrapolate the effects of high viscosity values (in an overestimated manner), these assumptions are still used. Figure S1A shows the results for $c \sim 68.5$ % wt/wt and $L = 2$m at the location $x_f \approx 25\%$ of the domain. From the area-averaged concentration profile, one can see that both models behave the same, as expected from equation (13). However, the axial velocity profile differs for both models. In the variable viscosity model, the maximum area-averaged velocity $\bar{U}$ is higher compared to the constant viscosity model. In addition, it has a wider non-zero range. Figure S1B shows the results for $L = 2$m at the location $x_f \approx 25\%$ of the domain for both models. An optimal sucrose concentration corresponding to a $J_{max}$ does exist for the variable viscosity model albeit at high values. In addition to having a higher optimal $c$ point, the profile of the variable viscosity model shows a tendency of having a higher range of efficiency compared to the constant viscosity model (for example, taking efficiency around 96%, the variable viscosity model have a range of $\approx$ 10 % wt/wt compared to $\approx$ 8 % wt/wt for the constant viscosity model). Since the assumptions might not hold at these high concentrations, revisions that accommodate non-Newtonian fluids and deviations from the Van't Hopf approximation must be used and are better kept for future work.

**A**

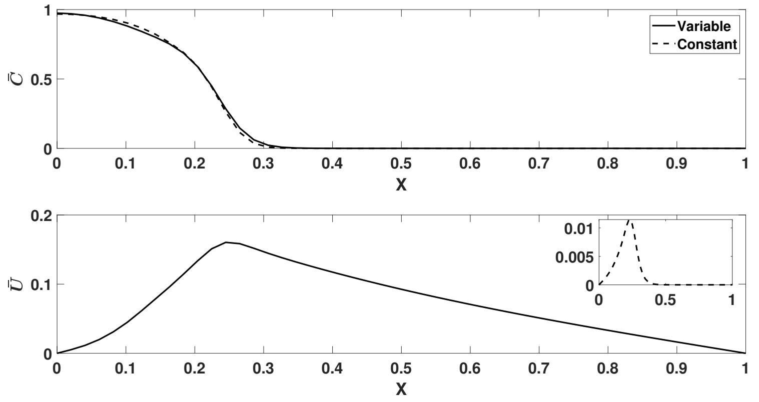


**B**

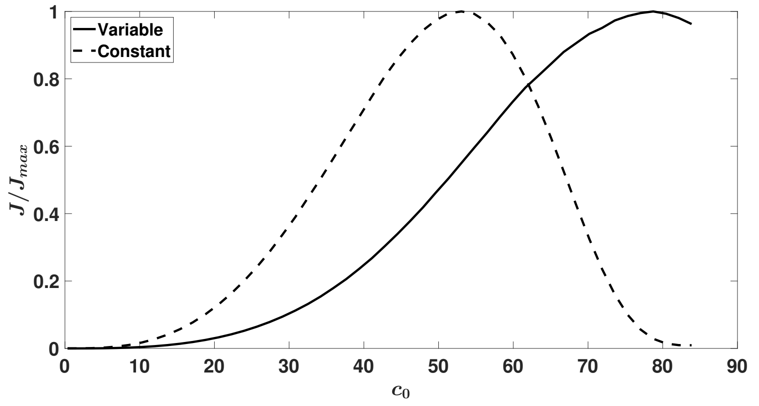


Figure S1: ($a$) Nondimensional area-averaged concentration $\bar{C}$ (upper figure) and axial velocity $\bar{U}$ (lower figure) profiles at the location $x_f \approx 0.25$ of the domain for $c_0 \sim 68.5$ % wt/wt and $L = 2$m. ($b$) Normalized flux $J/J_{max}$ as a function of initial concentration $c_0$. The variable viscosity model is denoted by a solid black line and the constant viscosity model is denoted by a solid dashed line.

### Materials and Methods S4 Thickness of the Unstirred Layer

In this section, the method to calculate the thickness of the 'unstirred layer' discussed in section 3.1 is presented. In osmotically driven flows, the bulk flow is driven by osmosis where the osmotic potential is calculated using the Van't hoff relation. This driving force is included in the model using a boundary condition that can be formulated from Darcy's law as discussed earlier. Since there is an influx of water, the concentration at the boundary decreases below the bulk concentration (i.e. concentration at the center of the tube). For this reason, the osmotic potential is lower than the one calculated using the bulk concentration as further discussed in Pedley (1983). In this work, the numerical model discussed in section Materials and Methods S2 takes into consideration the effects of this 'unstirred layer' because it solves the osmotic potential at the membrane wall using a zero concentration in water outside the membrane. However, because of the numerical approximation, there is an error that arises from spatially discretizing the domain and the effect of the layer might be underestimated, unless fully resolved by the spatial grid size (here $dr \sim 0.2\mu$m). To check if this layer is in fact resolved, the method discussed in Pohl et al. (1998) is used to first predict the thickness of the layer and then compare it to the spatial grid size. A summary of this method is presented as follow: a quadratic relation between the axial velocity and the radial coordinate $u(r) = a_1(1-r)^2$ is fitted to the simulations so as to solve for the constant $a_1$. This constant will then be used to solve for the thickness $\delta$ using equations 5 and 6 in Pohl et al. (1998). When this method was used for the example shown in section 2.2, the thickness of the layer was $\delta \sim 2.3\mu$m that is an order of magnitude larger than $dr$. It can be surmised that this layer was already resolved in the simulations here. Moreover, this unstirred layer has minor impact on the difference between the constant and variable viscosity model runs as it roughly cancels out when the solutions are substracted from each other.

**A**

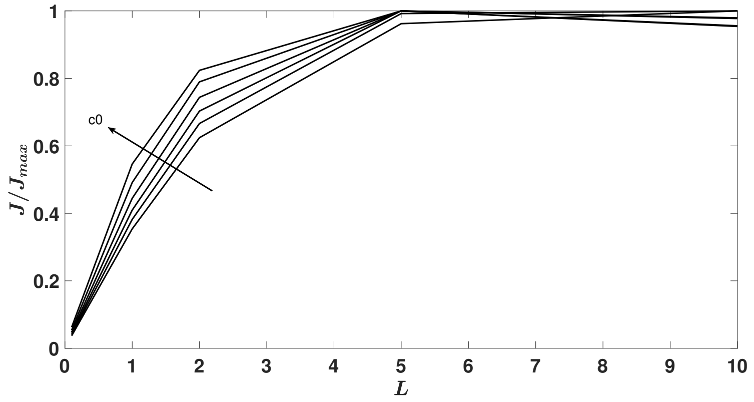


**B**

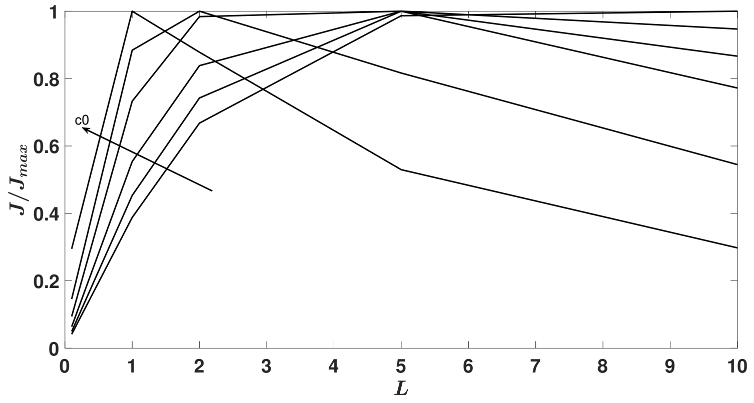


Figure S2: Normalized flux $J/J_{max}$ as a function of tube length $L$ for the $(a)$ variable viscosity model and $(b)$ constant viscosity model. Direction of the arrow indicates increasing loading concentration $c_0$ where $c_0 \sim 10$, 20, 30, 40, 50 and 60.
